# Association between Drusen Burden Determined by OCT and Genetic Risk in Early and Intermediate Age-Related Macular Degeneration

**DOI:** 10.1101/743633

**Authors:** Johanna M. Seddon, James Dossett, Rafael Widjajahakim, Bernard Rosner

## Abstract

**PURPOSE:** To determine associations between macular drusen parameters derived from an automatic optical coherence tomography (OCT) algorithm, age-related macular degeneration (AMD) stage and genetic variants.

**METHODS:** Eyes classified as early or intermediate AMD with OCT imaging and genetic data were selected (n=239 eyes). Drusen area and volume measurements were estimated using the Zeiss Cirrus advanced retinal pigment epithelium (RPE) analysis algorithm in a 5mm diameter (perifoveal) zone centered on the fovea. Associations between drusen measurements and common genetic variants in the complement and high density lipoprotein (HDL) lipid pathways and the *ARMS2* variant were calculated using generalized estimating equations and linear mixed models adjusting for age, sex, smoking, BMI, and education.

**RESULTS:** When compared to eyes with no measurable drusen, drusen area ≥ the median was independently associated with a higher number of risk alleles for *CFH* risk score, risk variants in *C3* and *ARMS2/HTRA1*. Similar results were obtained for drusen volume. When all genes were analyzed in the same model, only *CFH* score and *ARMS2/HTRA1* were associated with drusen measurements. HDL pathway genes were not significantly related to drusen parameters. Early and intermediate AMD stages were associated with OCT derived drusen area and volume.

**CONCLUSION:** Genetic variants in *CFH* and *ARMS2/HTRA1*, commonly associated with advanced AMD, were independently associated with higher drusen burden determined by OCT in eyes with early and intermediate AMD. The automatic RPE algorithm using OCT provides a quantitative classification of non-advanced AMD. Drusen morphology and other OCT-derived sub-phenotypes are biomarkers that could provide early anatomic endpoints for clinical trials.

## INTRODUCTION

Age-related macular degeneration (AMD) is a leading cause of vision loss and irreversible blindness in adults older than age 60.^1,2^ The etiology of AMD is multifactorial, and both behavioral and genetic risk factors contribute to personal risk.^2–4^ The genetic component of AMD risk is particularly well documented with regard to progression to advanced disease, including transitions to both advanced subtypes: geographic atrophy (GA) and neovascular disease (NV).^5–8^ Each advanced subtype is generally preceded by early and intermediate stages of disease that are primarily characterized by the formation of drusen between the retinal pigment epithelium (RPE) and Bruch’s membrane. The mechanisms by which an individual might develop GA or NV are not fully understood; however, it is clear that drusen development is predictive of progression to both forms of advanced disease.^9,10^ Management of the growing burden of advanced AMD remains a significant challenge. Therefore, identification of therapeutic targets for earlier, high risk stages of the disease is needed to lower the rate of progression to advanced stages and to preserve vision.

With advances in in vivo imaging, spectral-domain optical coherence tomography (SD-OCT) devices are capable of non-invasive visualization and discrimination of the retinal and choroidal layers. Dysfunction of the RPE, including area and volume abnormalities at locations where drusen develop, can be directly visualized and measured using OCT. Prior studies have indicated that OCT-derived drusen area and volume, and retinal features such as hyper-reflective foci and choroidal parameters, are informative in determining the likelihood of progression from early or intermediate to advanced AMD.^11–16^

However, knowledge about the association between OCT derived retina parameters and genetic variants related to AMD is sparse.^17–19^ We previously reported the effects of genes on progression to different stages of AMD.^20^ In that study, genes in the complement pathway were associated with a higher risk of progression from small and intermediate to large size drusen and large drusen to GA and NV. A novel association was also found between the *ABCA1* gene in the HDL pathway that reduced the risk of progression from normal to intermediate drusen and from intermediate to large drusen. Protective effects for transitions to advanced disease and larger drusen size were also observed in other pathways.^21,22^ These assessments provide a framework to evaluate the role of genetics as they relate to drusen measurements on OCT.

We evaluated the association between genetic risk and OCT drusen area and volume in a clinical cohort of patients with early and intermediate AMD. Understanding these relationships, in addition to other OCT parameters,^11–16,23^ may lead to clinical use of these measurements with identification of earlier high risk phenotypes and better stratification of risk of progression. Further characterization of drusen morphology on OCT may lead to the identification of disease biomarkers that could serve as anatomic endpoints for clinical trials. Our study aimed to evaluate the associations between a subset of genes implicated in risk of advanced AMD, and drusen area and volume measurements in eyes with clinically diagnosed early and intermediate AMD. (Seddon J, et al. AAO 2017; Abstract#30052798) (Widjajahakim R, et al. IOVS 2019; ARVO Abstract 3144907)

## METHODS

### Study Population and Classification of AMD Phenotypes

All participants were previously enrolled in our ongoing genetic and epidemiologic studies of AMD beginning in 1988.^24–27^ Participants were derived from clinic populations and nationwide referrals and were prospectively followed. The study protocol includes an ocular examination, fundus photography and OCT imaging, as well as interviews and blood sampling. This research adhered to the tenets of the Declaration of Helsinki and was approved by the institutional review board. Written informed consent was obtained for all participants.

AMD phenotypes were based on ocular examination and ocular imaging. Eyes were classified using the Clinical Age-Related Maculopathy Staging (CARMS) system by JMS as previously described.^28^ CARMS grades were defined as follows: grade 1 (no AMD, no drusen or only a few small drusen < 63 μm); grade 2 (early AMD, RPE irregularities and/or intermediate size drusen 63-124 μm); grade 3 (intermediate AMD, large drusen ≥ 125 μm); grade 4 (advanced dry AMD, or GA, including both central and non-central GA); and grade 5 (advanced exudative AMD, or NV, with choroidal neovascularization).

Risk factors were determined using a standardized questionnaire. Data for these analyses included demographic information-age, gender, education; anthropomorphic data-height and weight converted to body mass index; and the behavioral risk factor-smoking.

### Evaluation of Drusen Area and Volume Measurements

Beginning in 2007, OCT scans were obtained for the purpose of conducting studies to assess OCT parameters associated with various stages of AMD. In 2016, a retrospective review of Cirrus SD-OCT 4000 and 5000 scans was conducted among individuals with one or both eyes with stage 2 or 3 (Carl Zeiss Meditec Inc., Dublin, CA, USA). The OCT scanning protocols included high-definition 1 line scan, 5 line scan and macular cube over a 6×6 mm square, centered on the fovea. Drusen area and volume measurements were determined by the Cirrus Version 6.0 Advanced RPE Analysis algorithm, a fully automated algorithm available using the Cirrus SD-OCT instrument. This algorithm is based on an automatic determination of the RPE floor and measurement of the elevations of the RPE deformations. This yields an automatic quantitative assessment of RPE deformations and calculates measurements of area and volume that are highly reproducible.^29–31^ These measurements provide an estimate of drusen burden for a circle with 5 mm in diameter (perifoveal zone) centered at the fovea.^32^

Only eyes with early or intermediate AMD (CARMS grades 2 or 3) at the time of the OCT scan were selected for inclusion in the analyses. We selected participants with at least one OCT macula cube scan in at least one eye and with DNA genotyping data. Scans with signal strength less than 6 out of 10 were excluded similar to what has been previously reported.^15,33^ For individuals with multiple scans, the earliest, highest quality scan was evaluated.

### Genotyping and Genetic Data

Enrolled participants provided blood or saliva samples for DNA extraction according to a standard study protocol. Genotypes were determined using array-based and gene sequencing platforms as previously described.^24,25,34,35^ All single nucleotide polymorphisms (SNPs) had a high genotype call rate (>98%) and PLINK was used to perform all quality control steps.^36^

Common variants in genes previously associated with drusen and AMD in the complement pathway, HDL pathway, and the gene locus on chromosome 10q26 were selected given their consistent association with advanced disease or biologically plausible relationship to drusen formation. The genetic variants included complement factor H (*CFH*) Y402H rs1061170,^37^ *CFH* rs1410996,^34,38^ age-related maculopathy susceptibility 2/high-temperature requirement A serine peptidase 1 locus on chromosome 10q26 (*ARMS2* A69S/*HTRA1*) represented by SNP rs10490924,^39,40^ complement component 3 (*C3*) R102G rs2230199,^41^ and variants in the HDL pathway: hepatic lipase C (*LIPC*) rs10468017,^42^ adenosine tri-phosphate binding cassette transporter 1 (*ABCA1*) rs1883025,and cholesteryl ester transfer protein (*CETP* rs3764261.^20,42,43^

### Statistical Methods

A total of 239 eyes among 179 participants were included in these analyses. The distributions of demographic (age [<70, 70-79, ≥80], gender, education [≤high school,>high school], and behavioral [smoking and BMI], ocular [baseline AMD grade], and genetic factors) were evaluated for each area and volume measurement. Categorical comparisons were made between eyes with some drusen but < median vs eyes with no measurable drusen, and eyes with some drusen but ≥ median vs eyes with no measurable drusen in separate models for each outcome. Univariate associations between each genetic factor and the drusen measurements were evaluated by generalized linear models based on generalized estimating equations (GEE) using PROC GENMOD of SAS 9.4 with the individual eye as the unit of analysis, using a logistic link and a binomial distribution with a working independence correlation structure to account for the correlation between fellow eyes.^44^ We only considered eyes without advanced AMD since drusen morphology is altered by the presence of advanced AMD; thus some subjects contributed two eyes to the analyses, while other subjects contributed a single eye if for example the fellow eyes had advanced AMD. In addition, a multivariate model included all genetic factors in the same model. All models were adjusted for age, sex, smoking, BMI, and education. Two distinct outcomes of interest were assessed: drusen area and drusen volume in the perifoveal zone. Genetic variables were defined as having zero, one, or two risk or protective alleles. For the two *CFH* variants which convey different information about AMD risk (R^2^ = 0.44), we combined them into a risk score category from zero to four, consisting of the total number of alleles in the two variants combined. Odds ratios and 95% confidence intervals (CIs) were calculated per allele as estimates of effect size.

Since we identified consistent significant associations in categorical analyses between drusen measurements and risk genotypes for *CFH* risk score and *ARMS2*, we looked in more detail at mean area and volume represented as continuous variables for these two genes and assessed the independent effects of each gene on drusen measurements. For these analyses, AMD grade was included as an additional covariate. Associations between continuous drusen measurements and AMD stages and genotypes were evaluated by a linear mixed effects model using PROC MIXED of SAS 9.4 with the individual eye as the unit of analysis. This accounts for the inter-eye correlation in drusen area and volume between fellow eyes. The LSMEANS option of PROC MIXED was used to compute adjusted means of area and volume measurements for specific genotype categories and AMD grades. In addition, we used the LSMEANS option to compute adjusted means for area and volume of the *CFH* risk score, adjusted for age, sex, smoking, BMI, education, AMD grade, and *ARMS2* genotype (and similarly, LSMEANS for the *ARMS2* genotype was adjusted for *CFH* risk score).

We tested for significant interactions between *CFH* and *ARMS2* by first creating binary variables for each gene (*CFH* risk score: 3 or 4 alleles with reference = 0 to 2; and *ARMS2* GT and TT as 1 or 2 risk alleles with reference = GG as 0), and then creating the cross-product of the two binary variables. Finally we estimated Pearson correlation between area and volume using eyes as the unit of analysis.^45^ Two sided P values less than 0.05 were considered statistically significant. All analyses were conducted using SAS 9.4 (SAS Institute, Cary, NC, USA).

## RESULTS

Distributions of area and volume for different drusen categories are shown in **Supplementary Table 1**. More than 50% of eyes had some measurable drusen with a wide range in both area and volume among the subset of eyes with some measurable drusen. The correlation between area and volume for individual eyes was 0.84 indicating similarity between the two variables. Relationships between age, sex, smoking, BMI, and education, and drusen parameters comparing no measurable drusen vs < median drusen and no measurable drusen vs ≥ median drusen, were evaluated as shown in **Supplementary Table 2**. Older age was associated with greater drusen area and volume, and the other non-genetic factors were not significantly associated with these drusen measurements.

### Ocular Images of Drusen Measurements

Representative color fundus photographs and OCTs corresponding to different measurements using the advanced RPE analysis tool are shown in **Figure 1** for no measurable drusen, < median, and ≥ median drusen measurements.

**FIGURE 1.**
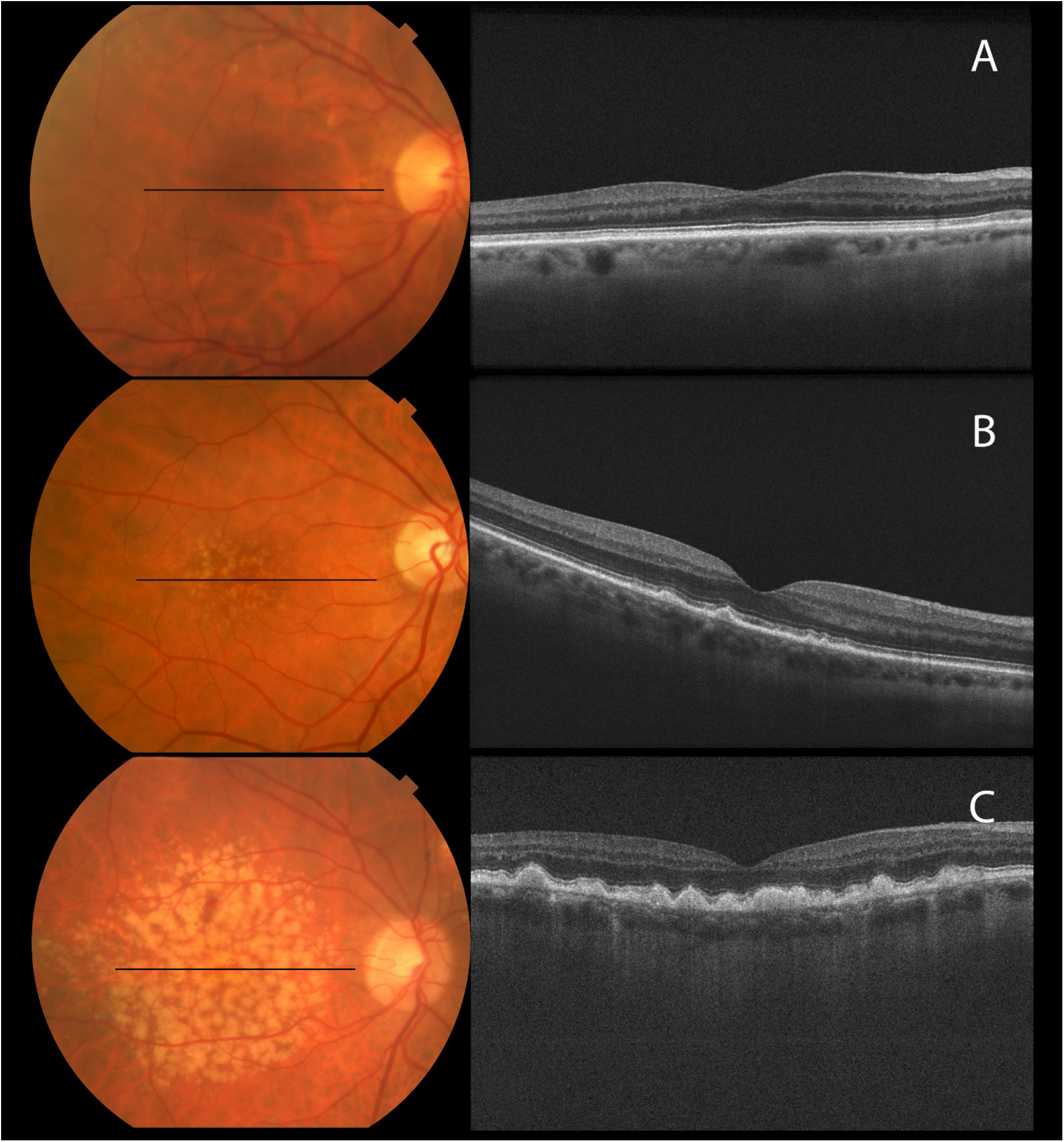
Representative color fundus photographs and OCTs corresponding to different measurements using the advanced RPE analysis tool are shown as A) no measurable drusen, B) drusen area and volume < median, and C) drusen area and volume ≥ median.

### Association between OCT Drusen Measurements and AMD Grade

The mean OCT derived drusen area was significantly associated with grade of AMD based on color photographs, and was higher in intermediate AMD eyes than in early AMD eyes, after adjusting for age, sex, smoking, BMI, and education (P = 0.008) (**Figure 2A**). Similarly, the mean drusen volume was higher in eyes with intermediate AMD compared with early AMD (P = 0.005) (**Figure 2B**).

**FIGURE 2.**
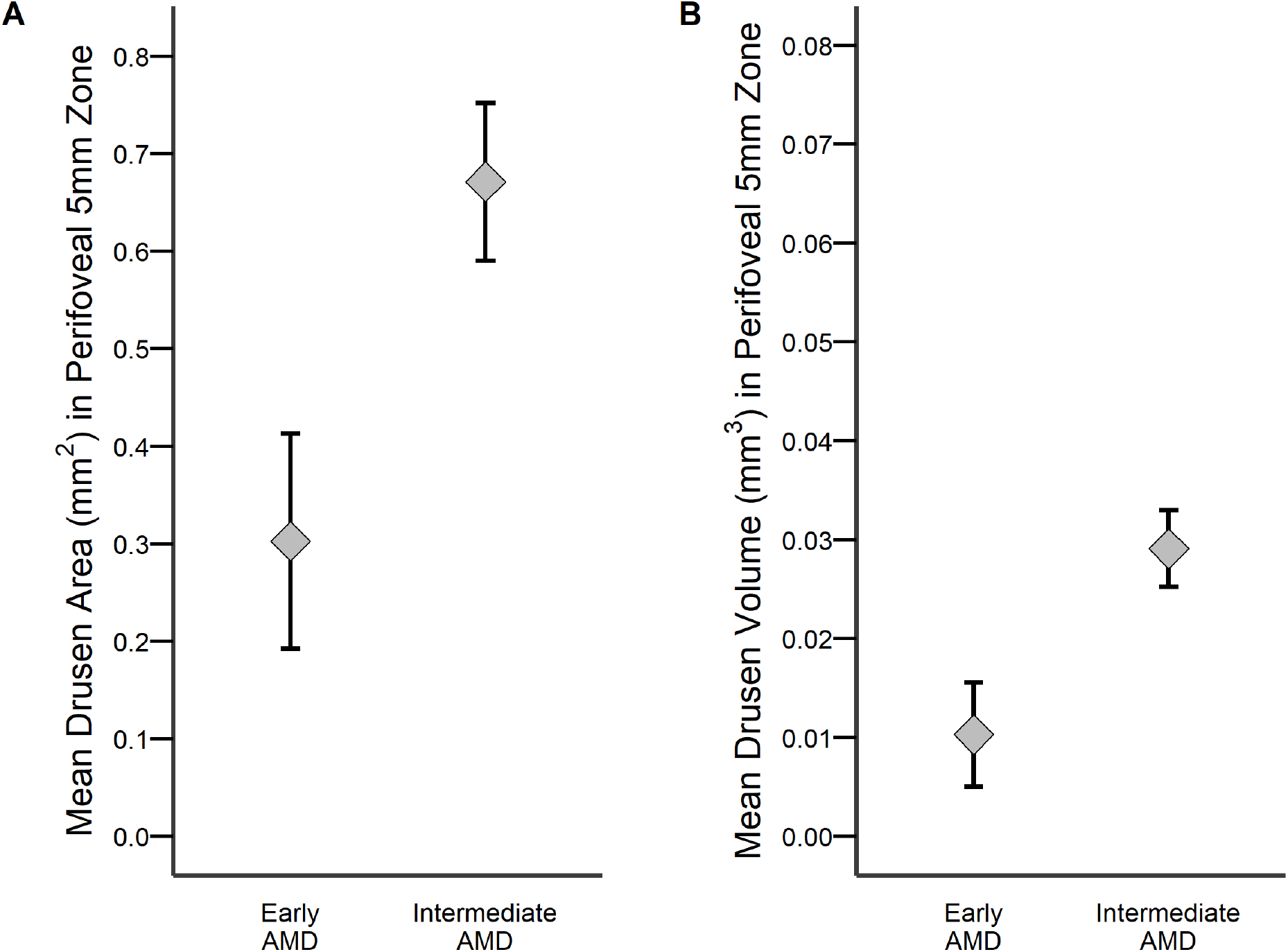
Associations between OCT-derived drusen measurements and early and intermediate stages of AMD, for A) drusen area (P=0.008), and B) drusen volume (P=0.005). The diamond represents the adjusted mean controlling for age, sex, smoking, BMI, and education. The vertical line represents plus or minus one standard error of the adjusted mean.

### Associations between Drusen Measurements and Each AMD Genetic Variant

A higher *CFH* score was associated with greater drusen area in the perifoveal 5mm zone controlling for age, sex, smoking, BMI, and education (**Table 1**). There was a significant association between drusen area ≥ median vs no-measurable drusen with OR = 1.79 (95% CI 1.18 – 2.71, P = 0.006) per category of *CFH* score. So the OR is 5.74 for category 3+ vs 0 (1.79^3^). For the variant in another complement pathway gene, *C3* R102G, there was a significant association between drusen area ≥ median vs no measurable drusen with OR = 1.80 (95% CI 1.07 – 3.03, P =0.03) per risk allele (G). A higher number of risk alleles for the variant in *ARMS2/HTRA1* was also associated with greater drusen area controlling for age, sex, smoking, BMI, and education. For the variant at this locus, there was a significant association between drusen area ≥ median vs no measurable drusen with OR = 2.76 (95% CI 1.57 – 4.86, P <0.001) per each risk allele (T). Genetic variants in the HDL pathway were not significantly associated with drusen area.

**TABLE 1.** Associations Between Drusen Area Measurements and Individual Genetic loci *

|  | No Drusen Measured<br>(N=111) | Drusen Area < median<br>(N=54) |  | Drusen Area ≥ median<br>(N=74) |  |
| --- | --- | --- | --- | --- | --- |
|  |  | OR<sup>†</sup> (95% CI) | P value<sup>‡</sup> | OR<sup>†</sup> (95% CI) | P value<sup>‡</sup> |
| *CFH* Score | 1.0 (ref) | 0.97 (0.65 - 1.47) | 0.90 | 1.79 (1.18 - 2.71) | 0.01 |
| *ARMS2/HTRA1*: rs10490924 | 1.0 (ref) | 1.29 (0.77 - 2.14) | 0.33 | 2.76 (1.57 - 4.86) | <0.001 |
| *C3* R102G: rs2230199 | 1.0 (ref) | 0.99 (0.57 - 1.72) | 0.98 | 1.80 (1.07 - 3.03) | 0.03 |
| *ABCA1*: rs1883025 | 1.0 (ref) | 0.66 (0.32 - 1.37) | 0.26 | 0.67 (0.35 - 1.31) | 0.24 |
| *LIPC*: rs10468017 | 1.0 (ref) | 1.14 (0.61 - 2.12) | 0.68 | 0.87 (0.51 - 1.50) | 0.62 |
| *CETP*: rs3764261 | 1.0 (ref) | 1.15 (0.67 - 1.98) | 0.62 | 1.49 (0.92 - 2.40) | 0.10 |

\* Each genetic variant assessed in a model, controlling for age, sex, education, BMI, and smoking.
<sup>†</sup> Odds ratio per risk or protective allele. *CFH* score categorized as 0,1,2,3-4 risk score categories.
<sup>‡</sup> Based on GEE using PROC GENMOD of SAS with the eye as the unit of the analysis using a logistic link and a binomial distribution with the working independence model to account for the inter-eye correlation. Separate analyses were performed i) comparing eyes with some drusen but less than the median vs eyes with no drusen, and ii) comparing eyes with some drusen but greater than or equal to the median vs eyes with no drusen.

Results for each genetic factor analyzed separately for association with the other drusen parameter, drusen volume, are shown in **Table 2**. Higher *CFH* score was associated with ≥ median drusen volume compared to no measurable drusen volume: OR = 1.73 (95% CI 1.13 – 2.64, P = 0.01) per category of *CFH* score. *C3* R102G was also related to drusen volume, with a significant association between number of risk alleles and drusen volume ≥ median compared to no drusen volume with OR = 1.92 (95% CI 1.09 – 3.41, P = 0.03) per risk allele. *ARMS2/HTRA1* risk was associated with greater drusen volume compared to no drusen volume (OR = 2.72, 95% CI 1.48-5.01, P = 0.001). The variant in the HDL gene, *CETP*, was borderline associated with higher drusen volume compared to no drusen volume with OR =1.63 (95% CI 0.99-2.68, P = 0.06).

**TABLE 2.** Associations Between Drusen Volume Measurements and Individual Genetic loci *

|  | No Drusen Measured<br>(N=96) | Drusen Volume < median<br>(N=66) |  | Drusen Volume ≥ median<br>(N=77) |  |
| --- | --- | --- | --- | --- | --- |
|  |  | OR<sup>†</sup> (95% CI) | P value<sup>‡</sup> | OR<sup>†</sup> (95% CI) | P value<sup>‡</sup> |
| *CFH* Score | 1.0 (ref) | 0.99 (0.66 - 1.48) | 0.96 | 1.73 (1.13 - 2.64) | 0.01 |
| *ARMS2/HTRA1*: rs10490924 | 1.0 (ref) | 1.29 (0.77 - 2.16) | 0.34 | 2.72 (1.48 - 5.01) | <0.001 |
| *C3* R102G: rs2230199 | 1.0 (ref) | 0.99 (0.55 - 1.79) | 0.98 | 1.92 (1.09 - 3.41) | 0.03 |
| *ABCA1*: rs1883025 | 1.0 (ref) | 0.69 (0.36 - 1.34) | 0.28 | 0.62 (0.32 - 1.22) | 0.17 |
| *LIPC*: rs10468017 | 1.0 (ref) | 0.94 (0.53 - 1.66) | 0.82 | 0.88 (0.50 - 1.53) | 0.65 |
| *CETP*: rs3764261 | 1.0 (ref) | 1.17 (0.69 - 1.99) | 0.56 | 1.63 (0.99 - 2.68) | 0.06 |

\* Each genetic variant assessed in a model, controlling for age, sex, education, BMI, and smoking.
<sup>†</sup> Odds ratio per risk or protective allele. *CFH* score categorized as 0,1,2,3-4 risk score categories.
<sup>‡</sup> Based on GEE using PROC GENMOD of SAS with the eye as the unit of the analysis using a logistic link and a binomial distribution with the working independence model to account for the inter-eye correlation. Separate analyses were performed i) comparing eyes with some drusen but less than the median vs eyes with no drusen, and ii) comparing eyes with some drusen but greater than or equal to the median vs eyes with no drusen.

### Multivariate Analyses of Associations between Drusen and Genetic Variants

When analyzing each genetic variable while controlling for all of the other variants, only two genetic variables, *CFH* risk score and the *ARMS2/HTRA1* variant remained independently associated with a significantly higher drusen area (≥ median compared to no measurable drusen) as shown in **Table 3**. For *CFH* score, the OR was 1.58 (95% CI 1.01-2.46, P= 0.04), and for the *ARMS2/HTRA1* variant, the OR was 2.45 (95% CI 1.35 – 4.45, P = 0.003).

**TABLE 3.** Multivariate Analyses of Associations Between Drusen Area and Genetic loci *

|  | No Drusen Measured<br>(N=111) | Drusen Area < median<br>(N=54) |  | Drusen Area ≥ median<br>(N=74) |  |
| --- | --- | --- | --- | --- | --- |
|  |  | OR<sup>†</sup> (95% CI) | P value<sup>‡</sup> | OR<sup>†</sup> (95% CI) | P value<sup>‡</sup> |
| *CFH* Score | 1.0 (ref) | 0.93 (0.61 - 1.41) | 0.73 | 1.58 (1.01 - 2.46) | 0.04 |
| *ARMS2/HTRA1*: rs10490924 | 1.0 (ref) | 1.24 (0.72 - 2.15) | 0.44 | 2.45 (1.35 - 4.45) | <0.001 |
| *C3* R102G: rs2230199 | 1.0 (ref) | 1.00 (0.55 - 1.82) | 0.99 | 1.57 (0.87 - 2.85) | 0.14 |
| *ABCA1*: rs1883025 | 1.0 (ref) | 0.65 (0.31 - 1.38) | 0.27 | 0.84 (0.41 - 1.73) | 0.65 |
| *LIPC*: rs10468017 | 1.0 (ref) | 1.14 (0.61 - 2.12) | 0.69 | 0.76 (0.43 - 1.36) | 0.36 |
| *CETP*: rs3764261 | 1.0 (ref) | 1.17 (0.67 - 2.04) | 0.57 | 1.28 (0.78 - 2.12) | 0.33 |

\* Each genetic variant assessed in a model, controlling for age, sex, education, BMI, smoking, and other genetic variants.
<sup>†</sup> Odds ratio per risk or protective allele. *CFH* score categorized as 0,1,2,3-4 risk score categories.
<sup>‡</sup> Based on GEE using PROC GENMOD of SAS with the eye as the unit of the analysis using a logistic link and a binomial distribution with the working independence model to account for the inter-eye correlation. Separate analyses were performed i) comparing eyes with some drusen but less than the median vs eyes with no drusen, and ii) comparing eyes with some drusen but greater than or equal to the median vs eyes with no drusen.

Multivariate associations between genes and drusen volume controlling for all genes are shown in **Table 4**. Comparing drusen volume ≥ median to no measurable drusen, *CFH* score had an OR of 1.54 (95% CI 0.97-2.45, P = 0.07). *ARMS2/HTRA1* remained significant with an OR = 2.49 (95% CI 1.29 – 4.80, P = <0.001). The *C3* and *CETP* variants were not associated with higher drusen volume (P=0.09).

**TABLE 4.** Multivariate Analyses of Associations Between Drusen Volume and Genetic loci *

|  | No Drusen Measured<br>(N=96) | Drusen Volume < median<br>(N=66) |  | Drusen Volume ≥ median<br>(N=77) |  |
| --- | --- | --- | --- | --- | --- |
|  |  | OR<sup>†</sup> (95% CI) | P value<sup>‡</sup> | OR<sup>†</sup> (95% CI) | P value<sup>‡</sup> |
| *CFH* Score | 1.0 (ref) | 0.95 (0.64 - 1.42) | 0.81 | 1.54 (0.97 - 2.45) | 0.07 |
| *ARMS2/HTRA1*: rs10490924 | 1.0 (ref) | 1.27 (0.74 - 2.18) | 0.39 | 2.49 (1.29 - 4.80) | 0.01 |
| *C3* R102G: rs2230199 | 1.0 (ref) | 1.00 (0.53 - 1.90) | 1.00 | 1.78 (0.92 - 3.45) | 0.09 |
| *ABCA1*: rs1883025 | 1.0 (ref) | 0.69 (0.36 - 1.35) | 0.28 | 0.76 (0.37 - 1.57) | 0.45 |
| *LIPC*: rs10468017 | 1.0 (ref) | 0.91 (0.51 - 1.62) | 0.75 | 0.73 (0.41 - 1.32) | 0.30 |
| *CETP*: rs3764261 | 1.0 (ref) | 1.19 (0.69 - 2.04) | 0.53 | 1.54 (0.93 - 2.54) | 0.09 |

\* Each genetic variant assessed in a model, controlling for age, sex, education, BMI, smoking, and other genetic variants.
<sup>†</sup> Odds ratio per risk or protective allele. *CFH* score categorized as 0,1,2,3-4 risk score categories.
<sup>‡</sup> Based on GEE using PROC GENMOD of SAS with the eye as the unit of the analysis using a logistic link and a binomial distribution with the working independence model to account for the inter-eye correlation. Separate analyses were performed i) comparing eyes with some drusen but less than the median vs eyes with no drusen, and ii) comparing eyes with some drusen but greater than or equal to the median vs eyes with no drusen.

### Independent Associations between Drusen Measurements and CFH Score and ARMS2

As shown in **Table 5**, both mean perifoveal drusen area and volume increased as the *CFH* score increased, and the trends for increasing drusen burden as the *CFH* score increased were significant after adjusting for age, sex, smoking, BMI, education, and also AMD grades (P trend = 0.004 for area and P trend= 0.002 for volume). Carriers of 2 risk alleles versus 0-1 had higher drusen area (P=0.03) and drusen volume (P=0.04), and carriers of 3-4 risk alleles also had significantly higher drusen area (P= <0.001) and volume (P=<0.001) compared with having 0-1 risk alleles. Similar comparisons were assessed for the *ARMS2/HTRA1* variant: mean drusen area and volume increased as the number of risk alleles increased, after adjustment for other variables (P trend <0.001 for both drusen area and volume). Carrying two risk alleles versus none was significantly related to higher drusen area and volume (P= 0.008 and 0.004, respectively). These associations between OCT-derived perifoveal drusen measurements and genetic factors are also depicted in **Figure 3**.

**TABLE 5.** Associations Between Drusen Area and Volume and *CFH* and *ARMS2/HTRA1*

| No. of Risk Alleles | Univariate Gene Analysis* |  |  |  |  | Bivariate Gene Analysis† |  |  |  |
| --- | --- | --- | --- | --- | --- | --- | --- | --- | --- |
|  | n<br>(eyes) | Drusen Area<br>Mean<br>(mm<sup>2</sup>±SE) | P<br>Value | Drusen Volume<br>Mean<br>(mm<sup>3</sup>±SE) | P<br>Value | Drusen Area<br>Mean<br>(mm<sup>2</sup>±SE) | P<br>Value | Drusen Volume<br>Mean<br>(mm<sup>3</sup>±SE) | P<br>Value |
| ***CFH* Risk Score‡** |  |  |  |  |  |  |  |  |  |
| 0-1 | 39 | 0.07 ± 0.16 | Ref | 0.0002 ± 0.008 | Ref | 0.25 ± 0.17 | Ref | 0.007 ± 0.008 | Ref |
| 2 | 103 | 0.49 ± 0.10 | 0.03 | 0.019 ± 0.005 | 0.04 | 0.57 ± 0.11 | 0.09 | 0.022 ± 0.005 | 0.11 |
| 3-4 | 97 | 0.66 ± 0.10 | <0.001 | 0.029 ± 0.005 | <0.001 | 0.74 ± 0.11 | 0.01 | 0.032 ± 0.005 | 0.006 |
| P (trend) |  |  | 0.004 |  | 0.002 |  | 0.01§ |  | 0.005§ |
| ***ARMS2/HTRA1*:**<br>**rs10490924** |  |  |  |  |  |  |  |  |  |
| 0 | 105 | 0.194 ± 0.10 | Ref | 0.007 ± 0.005 | Ref | 0.15 ± 0.098 | Ref | 0.005 ± 0.0047 | Ref |
| 1 | 104 | 0.677 ± 0.10 | <0.001 | 0.028 ± 0.005 | 0.002 | 0.60 ± 0.10 | 0.001 | 0.024 ± 0.005 | 0.004 |
| 2 | 30 | 0.898 ± 0.19 | 0.008 | 0.036 ± 0.009 | 0.004 | 0.80 ± 0.19 | 0.002 | 0.031 ± 0.0089 | 0.009 |
| P (trend) |  |  | <0.001 |  | <0.001 |  | <0.001\|\| |  | 0.001\|\| |

\* Analyses based on continuous area and volume, adjusting for age, sex, BMI, smoking, AMD grade, and education using the LSMEANS option of PROC MIXED of SAS. Each genetic factor was analyzed separately.
† Analyses for *CFH* score are based on continuous area and volume, adjusting for age, sex, BMI, smoking, education, AMD grade, and *ARMS2/HTRA1* using the LSMEANS option of PROC MIXED of SAS. Analyses for *ARMS2/HTRA1* were performed similarly, adjusting for *CFH* score.
‡ No. of risk alleles for rs1410996 and rs1061170 combined.
§ The P trend assessing risk per 1 category increase in *CFH* score was obtained after adjusting for categories of the *ARMS2/HTRA1* and other risk factors.
|| The P trend assessing risk per 1 category increase in *ARMS2/HTRA1* was obtained after adjusting for categories of the *CFH* score and other risk factors.

**FIGURE 3.**
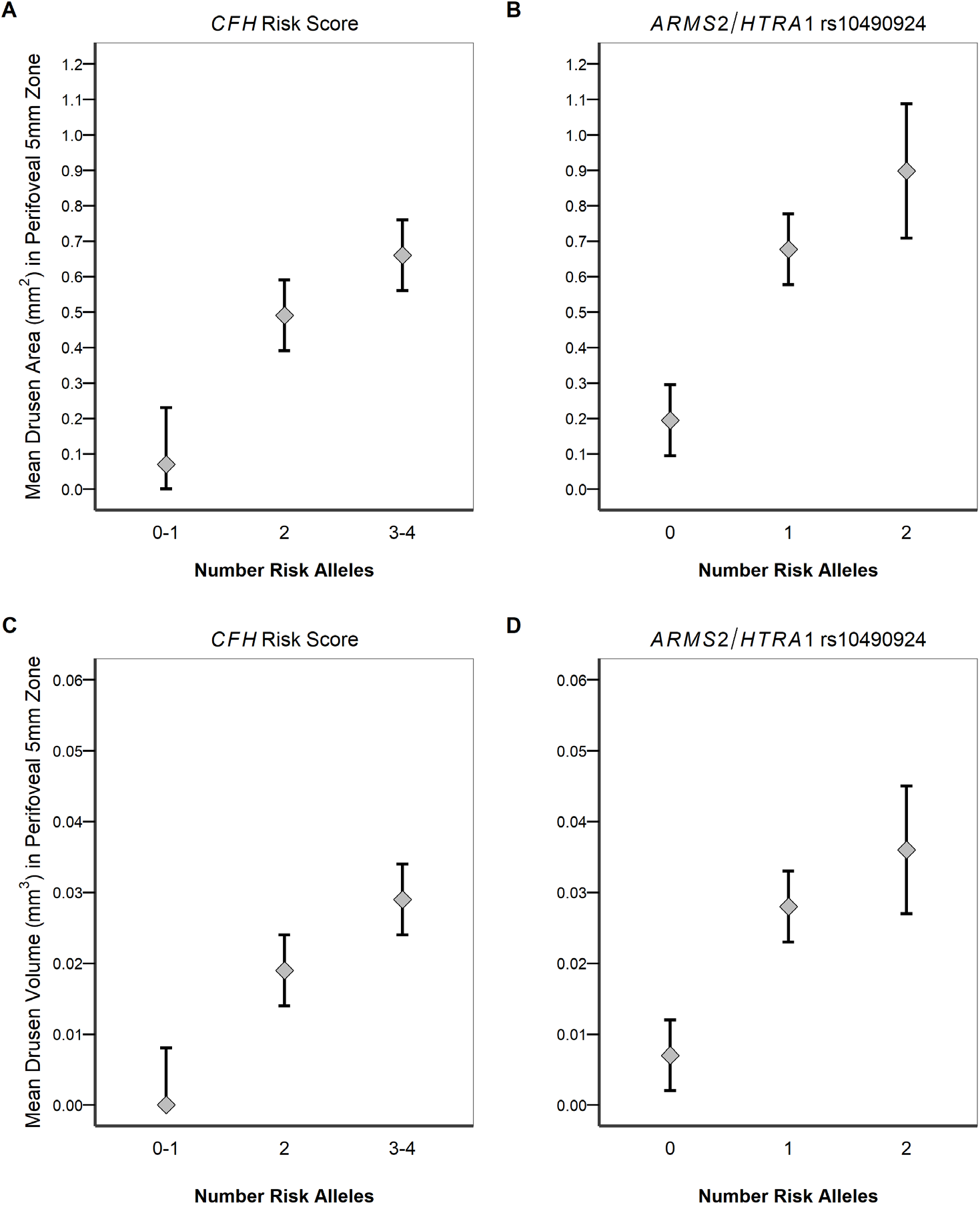
Independent associations between OCT-derived perifoveal drusen measurements and number of risk alleles for drusen area for A) *CFH* score and B) *ARMS2/HTRA1* SNP rs10490924 (P trend= 0.004 and <0.001 respectively), and drusen volume for C) *CFH* score and D) *ARMS2/HTRA1* SNP rs10490924 (P trend = 0.002 and <0.001 respectively). The diamond represents the adjusted mean controlling for age, sex, smoking, BMI, education, and AMD grade. The vertical line represents plus or minus one standard error of the adjusted mean.

When both genes were adjusted for each other (bivariate analyses shown in **Table 5**), the associations between drusen measurements and *CFH* score were somewhat reduced. However, the trend for higher drusen area and volume with higher score remained significant (P trend= 0.01 for area; P trend= 0.005 for volume). When the *ARMS2/HTRA1* genetic variant was adjusted for the *CFH* score, results were essentially unchanged from the univariate analysis as above, and trends for increasing drusen area and volume with increasing number of *ARMS2/HTRA1* risk alleles were significant (P trend= <0.001 and 0.001 for area and volume respectively).

All tests of interaction between *CFH* score and *ARMS2/HTRA1* were not significant. When comparing drusen area or volume measurements ≥ median vs no measurable drusen, the P values were 0.38 for drusen area and 0.33 for drusen volume. (Data not shown)

## DISCUSSION

We evaluated associations between measurements of drusen area and volume and risk and protective alleles known to be related to advanced AMD, in patients with early and intermediate AMD. Drusen area and volume measurements were estimated using the advanced RPE analysis algorithm from Zeiss Cirrus OCT. These measurements were greater for eyes classified as having intermediate AMD compared with early AMD. In addition, among eyes with the same AMD grade (early or intermediate), genetic risk was associated with a higher drusen burden. *CFH* score and *ARMS2/HTRA1* were independently associated with perifoveal drusen area and volume compared with no measurable drusen.

Interestingly, genes in the HDL pathway were not significantly associated with drusen area and volume. Significant associations between drusen parameters and the *C3* variant were seen in univariate analyses but not in the multivariate analyses.

When comparing the adjusted means of the drusen area and volume for *CFH* score and *ARMS2/HTRA1*, we found significant trends for increasing drusen measurements with increasing number of risk alleles for each variable. The trends persisted when both genes were adjusted for each other as well as when the baseline AMD grade was included as a covariate in the analysis. There was no interaction between these genetic factors when assessing their association with drusen parameters. These two genetic variants contributed independently to drusen burden.

Prior studies investigating the role of genetics and drusen assessments based on OCT are limited and inconsistent. Chavali et al. studied an Amish population with early AMD and found an association between drusen and risk alleles in *CFH* rs12038333 and SYN3 rs5749482, but not *ARMS2*.^18^ Authors reported that the population had a high rate of homozygous risk allele of *SYN3*, and suggested that this population has a unique genetic background. Oeverhaus et al. found that among 85 patients in Germany, individuals homozygous for *CFH* Y402H or *ARMS2* A69S had larger amounts of drusen and different types of drusen based on manual OCT measurements, although measurements of area and volume of drusen based on an automatic OCT algorithm were not assessed.^46^ A study of change in drusen volume over one year among 30 patients in Florida who participated in a study of eculizamab showed an association with *CFH* rs1061170 but not *ARMS2*.^19^ Drusen volume based on OCT was assessed in the Singapore Eye Disease program and only *ARMS2* was associated with drusen volume, although the *CFH* SNPs in our risk score and drusen area were not evaluated.^47^

The value in obtaining and following these OCT drusen measurements over time has been suggested. Folgar et al. showed that greater baseline OCT drusen volume was associated with increased risk of progression to NV over two years.^14^ Sleiman et al. demonstrated that OCT based drusen measurements were associated with appearance of geographic atrophy on color photographs over 4 years.^48^ Schmidt-Erfurth et al found that drusen area was an important quatitative feature for progression.^16^ Higher drusen volume was related to advanced AMD in a retrospective review by Lei et al, but other OCT parameters were more strongly related in their cases.^23^ A combination of drusen parameters along with other OCT derived parameters such as hyperreflective foci and retinal thickness may be the most informative and predictive of progression.^15,16,23^

The association between advanced AMD and the *CFH* Y402H variant as well as the intronic SNP in this gene, *CFH* rs1410996, have been well documented ^6,20,34,38,49–51^ The biologic mechanisms underlying their effect on drusen accumulation, however, have not been fully explored. In our previous analyses of these variants regarding progression, the effect of rs1410996 was stronger and the effect of Y402H was reduced when both were in the same model, suggesting that this variant may be more strongly associated with AMD. *CFH* negatively regulates the alternate complement pathway and dampens the excessive C3 convertase activated by either immune complex deposition or C3 convertase activation from pathogens or damaged cell surfaces. *CFH* risk variants are functionally less efficient at dampening down this response leading to heightened complement activity which can lead to AMD related pathology.

It should be noted that SNPs in the genes *ARMS2* and *HTRA1* at the chromosome 10q26 locus are in very high linkage disequilibrium and functional studies are needed to determine which gene products lead to AMD pathology.^40^ The function of the ARMS2 protein in humans and the consequences of the A69S variant have been explored but have not been confirmed. One study found that *ARMS2* was expressed in human monocytes and microglia cells and facilitated removal of cellular debris by local complement activation; the A69S variant resulted in ARMS2 deficiency, possibly impairing the removal of cellular debris at Bruch’s membrane, leading to the development of drusen.^52^ On the other hand, another study reported that ARMS2 mRNA and protein are expressed at extremely low levels in eye tissues, and presented support for *HTRA1* as the risk factor for AMD. ^53^

We found that the OCT algorithm yielded drusen measurements that were highly associated with the manual CARMS grading. This advanced RPE analysis algorithm or a similar method may be useful for ophthalmologists or reading centers to categorize the stage of AMD more efficiently, with higher precision and with the ability to quantify the measurements, compared with clinical observation or photography. Higher resolution OCT may reduce variability in OCT grading and enhance the application of the RPE analysis algorithm. Future studies will continue to provide more information about the impact of genetic variants on drusen morphology.

### Strengths

Strengths of this study include the assessment of plausible genetic pathways, the inclusion of high quality scans, and use of a commercially available means of estimating drusen measurements that is incorporated into Cirrus SD-OCT instruments. Automated RPE analysis has been shown to be consistent and reproducible, especially between repeated scans.^31,54^ We also conducted eye specific analyses adjusting for correlation between eyes, analyzed both drusen area and volume, and controlled for demographic and behavioral risk factors associated with AMD.

### Limitations

Some of the AMD-associated genes did not show an association with drusen area or volume in eyes with early and intermediate AMD. This may be due to lack of an association or insufficient power secondary to a limited number of eyes in the homozygous risk or protective genotype category in some of these genes. We did not find a significant association between OCT-derived drusen measurement and the HDL pathway genes (*ABCA1, LIPC*, and *CETP*), although effect estimates were in the direction previously seen in a prospective analysis.^20^ Of note, the same genes (*CFH* and *ARMS2/HTRA1*) significantly related to the transition from early to intermediate AMD in Yu et al^20^ based on color photographs were also associated with larger OCT-derived drusen measurements in our analyses as seen in Tables 1 to 5 of the current paper.

We observed that some small drusen seen on color photographs were classified as having no measurable drusen by the RPE algorithm on OCT. This may due to the basis of the algorithm that has a threshold of 10 pixels elevation before the measurements can be calculated.^30^ The developers of the algorithm used this to prevent a false positive RPE elevation in the OCT due to noise. Furthermore, drusen outside the perifoveal zone were not measured since they were located outside the scanned or analyzed zone. Since the same algorithm for measuring drusen parameters was applied to all data uniformly, the relative rankings of the area and volume measurements were internally valid.

### Summary

In summary, this study determined that risk alleles in AMD-associated genes, *CFH* and *ARMS2/HTRA1*, were independently associated with greater drusen area and volume based on automated measurements of RPE using the Zeiss SD-OCT algorithm. The associations between these genes and drusen remained significant after controlling for all other genes as well as the non-genetic covariates. The *C3* variant was associated with drusen area and volume only in univariate analyses. Results confirm some findings in the few previous studies in different populations, but also contribute new information and analyses to support the involvement of both *CFH* and *ARMS2/HTRA1* genes in drusen development.

As our understanding of the genetic determinants of AMD becomes clearer, it will be important to understand the specific patho-biologic consequences of these genetic variants. SD-OCT can quantify variation in morphology in a way that is accurate and reproducible. Other AMD phenotypes detectable on OCT including subfoveal fluid, retinal and choroidal thickness, retinal hyperreflective foci, other drusen characteristics such as hyper-reflectivity and homogeneity, are opportunities for exploration in the context of subtyping AMD and assessing risk of progression,^15,16,23,48^ as well as defining the associations with genetic variants. Understanding the relationship between disease severity, morphologic features, and genetic factors has the potential to enhance individualized patient management and treatment.

Genetic risk is associated with a higher drusen burden assessed by OCT in eyes with early and intermediate stages of AMD. Further characterization of drusen and retinal and choroidal morphology may lead to the identification of biomarkers that could serve as early anatomic endpoints for clinical trials.

## ACKNOWLEDGEMENTS

*Funding/Support:* NIH Grants R01 EY011309 and R01 EY022445; Massachusetts Lions Eye Research Fund. Inc; International Retina Foundation; American Macular Degeneration Foundation.

*Financial Disclosures:* JMS: Gemini Therapeutics, Inc. Senior Medical Advisor; THEA Pharmaceuticals, Scientific Advisory Board; JD: none, RW: none, BR: none.

All authors attest that they meet the current ICMJE criteria for authorship

