## Supplemental Tables for "Association between Drusen Burden Determined by OCT and Genetic Risk in Early and Intermediate Age-Related Macular Degeneration"

**SUPPLEMENTAL TABLE 1.** Distribution of Drusen Measurements

| | N (% eyes) | Mean (SD)[*] [†] | Median[*] | IQR[*] | Correlation[‡] |
|---|---|---|---|---|---|
| **Area** | | | | | 0.84 (0.72 - 0.92) |
| no drusen measured | 111 ( 46.4 ) | - | - | - | |
| some measurable drusen | 128 ( 53.6 ) | 1.01 ± 1.22 | 0.70 | 0.25 - 1.10 | |
| <median | 54 ( 22.6 ) | 0.22 ± 0.12 | 0.20 | 0.10 - 0.30 | |
| ≥median | 74 ( 31.0 ) | 1.59 ± 1.33 | 1.00 | 0.80 - 1.90 | |
| **Volume** | | | | | |
| no drusen measured | 96 ( 40.2 ) | - | - | - | |
| some measurable drusen | 143 ( 59.8 ) | 0.038 ± 0.06 | 0.019 | 0.003 - 0.039 | |
| <median | 66 ( 27.6 ) | 0.005 ± 0.004 | 0.003 | 0.001 - 0.009 | |
| ≥median | 77 ( 32.2 ) | 0.065 ± 0.07 | 0.038 | 0.024 - 0.076 | |

[*] Among eyes with some measurable drusen.
[†] Unit for area is $mm^2$ and for volume is $mm^3$.
[‡] Pearson correlation between area and volume for individual eyes adjusting for correlation between fellow eyes.[45]

**SUPPLEMENTAL TABLE 2.** Associations Between Drusen Area and Volume Measurements and Non Genetic Factors

| | No Drusen Measured | Drusen < Median | P Value* | Drusen ≥ Median | P Value† |
|---|---|---|---|---|---|
| **Area** | | | | | |
| **Age (years)** | | | | | |
| < 70 | 40 (36) | 17 (31) | Ref | 14 (19) | Ref |
| 70-79 | 33 (30) | 16 (29) | 0.82 | 27 (36) | 0.07 |
| ≥ 80 | 38 (34) | 21 (39) | 0.53 | 33 (45) | 0.02 |
| **Sex** | | | | | |
| Male | 34 (31) | 21 (39) | Ref | 25 (34) | Ref |
| Female | 77 (69) | 33 (61) | 0.48 | 49 (66) | 0.78 |
| **Education** | | | | | |
| ≤ High school | 38 (35) | 15 (28) | Ref | 23 (31) | Ref |
| > High school | 72 (65) | 39 (72) | 0.64 | 51 (69) | 0.95 |
| **BMI** | | | | | |
| < 25 | 43 (39) | 23 (43) | Ref | 22 (30) | Ref |
| ≥ 25 | 68 (61) | 31 (57) | 0.45 | 52 (70) | 0.55 |
| **Smoking** | | | | | |
| Never | 41 (38) | 22 (41) | Ref | 31 (42) | Ref |
| Ever | 68 (62) | 32 (59) | 0.44 | 43 (58) | 0.55 |
| **Volume** | | | | | |
| **Age (years)** | | | | | |
| < 70 | 38 (40) | 18 (27) | Ref | 15 (19) | Ref |
| 70-79 | 26 (27) | 24 (36) | 0.12 | 26 (34) | 0.06 |
| ≥ 80 | 32 (33) | 24 (36) | 0.34 | 36 (47) | 0.02 |
| **Sex** | | | | | |
| Male | 28 (29) | 25 (38) | Ref | 27 (35) | Ref |
| Female | 68 (71) | 41 (62) | 0.42 | 50 (65) | 0.88 |
| **Education** | | | | | |
| ≤ High school | 32 (34) | 21 (32) | Ref | 23 (30) | Ref |
| > High school | 63 (66) | 45 (68) | 0.89 | 54 (70) | 0.98 |
| **BMI** | | | | | |
| < 25 | 39 (41) | 26 (39) | Ref | 23 (30) | Ref |
| ≥ 25 | 57 (59) | 40 (61) | 0.90 | 54 (70) | 0.41 |
| **Smoking** | | | | | |
| Never | 37 (39) | 23 (35) | Ref | 34 (44) | Ref |
| Ever | 57 (61) | 43 (65) | 0.76 | 43 (56) | 0.51 |

* P value was based on GEE using PROC GENMOD of SAS with the eye as the unit of the analysis using a logistic link and a binomial distribution with the working independence model to account for the inter-eye correlation for measurable drusen vs drusen < median.
† Similar analysis for no measurable drusen vs drusen ≥ median
